# Investigate the relevance of major signaling pathways in cancer survival using a biologically meaningful deep learning model

**DOI:** 10.1101/2020.04.13.039487

**Authors:** Jiarui Feng, Heming Zhang, Fuhai Li

## Abstract

Survival analysis and prediction are important in cancer studies. In addition to the Cox proportional hazards model, recently deep learning models have been proposed to integrate the multi-omics data for survival prediction. Cancer signaling pathways are important and interpretable concepts that define the signaling cascades regulating cancer development and drug resistance. Thus, it is interesting and important to investigate the relevance to patients’ survival of individual signaling pathways. In this exploratory study, we propose to investigate the relevance and difference of a small set of core cancer signaling pathways in the survival analysis of cancer patients. Specifically, we built a biologically meaningful and simplified deep neural network, DeepSigSurvNet, for survival prediction. In the model, the gene expression and copy number data of 1648 genes from 46 major signaling pathways are used. We applied the model on 4 types of cancer and investigated the relevance and difference of the 46 signaling pathways among the 4 types of cancer. Interestingly, the interpretable analysis identified the distinct patterns of these signaling pathways, which are helpful to understand the relevance of the signaling pathways in terms of their association with cancer survival time. These highly relevant signaling pathways can be novel targets, combined with other essential signaling pathways inhibitors, for drug and drug combination prediction to improve cancer patients’ survival time.

## Introduction

Survival analysis is important in cancer prognosis based on the clinical factors, e.g., age, gender, race, stage. Moreover, it is important to identify and understand essential biomarkers with the availability of large-scale genomics data, e.g., gene expression and copy number variation, in addition to these clinical factors. The cox proportional hazards model (Cox PH) model^1^ is the classic model for the survival analysis. Recently, the deep learning models have been widely used in image analysis^2,3^, medical informatics data analysis^4^, nature language process (NLP)^5^, and showed significantly improved performance than traditional machine learning models. Deep learning models have been developed for survival analysis.

Compared with the Cox PH model, the deep learning models showed improved prediction accuracy by integrating a large number of genomics features. For example, for the liver cancer subtyping and survival analysis^6^, the auto-encoder model was first employed to reduce the dimension of feature space, considering the large-number of genomics features, e.g., gene expression, miRNA, Methylation. The important features (non-linear combinations of raw genomics features) were identified using the Cox PH model^1^ for the clustering analysis to identify sub-groups with distinct survival outcomes. Then the analysis of variance (ANOVA), based on the clustering results, was applied on the raw genomics features was further used to identify the important genes. However, the auto-encoder model itself was not analyzed to identify the important raw genomics features in a non-linear perspective. In the Cox-nnet model, the RNA-seq data of TCGA samples was used as the input a deep neural network to predict the survival time. To identify the potential associated signaling pathways of hidden nodes, the Pearson’s correlation values between the expression of individual genes and the output of the given hidden nodes were calculated to identify the most linearly correlated genes. Then gene set enrichment analysis (GSEA)^7^ was employed to link the hidden nodes with the enriched signaling pathways. Moreover, the Survival Convolutional Neural Networks (SCNN)^8^ was developed to predict the survival using histologic images of cancer patients. Then heat map visualization of SCNN model output of regions of interest (image patches) was overlaid on the image to indicate the important regions in the images correlated with survival outcome.

In cancer studies, many dysfunctional signaling pathways are identified^9^, which play important roles in tumor development and drug response. In this study, we aim to investigate the relevance of these signaling pathways in the context of survival outcome prediction using a biologically meaningful and simplified deep learning model, DeepSigSurvNet. Specifically, only signaling pathways (46 pathways) were collected from KEGG^10^ signaling database. The gene expression and copy number data of 1648 genes from the 46 major signaling pathways are from 4 types of cancer: breast invasive carcinoma (BRCA), lung adenocarcinoma (LUAD), glioblastoma multiforme (GBM), and skin cutaneous melanoma (SKCM). The model was evaluated using the c-index.

Interestingly, the interpretable analysis using the Layer-wise relevance propagation (LRP)^11^ approach identified distinct probability density distribution patterns of these signaling pathways, which are helpful to understand the relevance of the signaling pathways in terms of the association with cancer patients’ survival. The important signaling pathways can be novel targets for drug and drug combination prediction to improve cancer patients’ survival time. In the following sections, the materials and methods, results and discussions are presented.

## Materials and Methodology

### RNA-seq and Copy number data of 4 types of cancer

From the UCSC Xena data server, the mean-normalized log2 scaled RSEM^12^ values (per gene) across all TCGA cohorts (HiSeqV2_PANCAN dataset) and integer copy number data (per gene) from GISTIC2 analysis were downloaded for 4 types of cancer: breast invasive carcinoma (BRCA), lung adenocarcinoma (LUAD), glioblastoma multiforme (GBM), and skin cutaneous melanoma (SKCM). The phenotype (clinical) data, including survival time, age, gender, stage, of the cancer samples are also available from the Xena data server. Table I shows the number of cancer samples, dataset and URLs to download these datasets. For the prediction purpose, cancer patients with survival time greater than 3000 days are not included.

**Table I:**
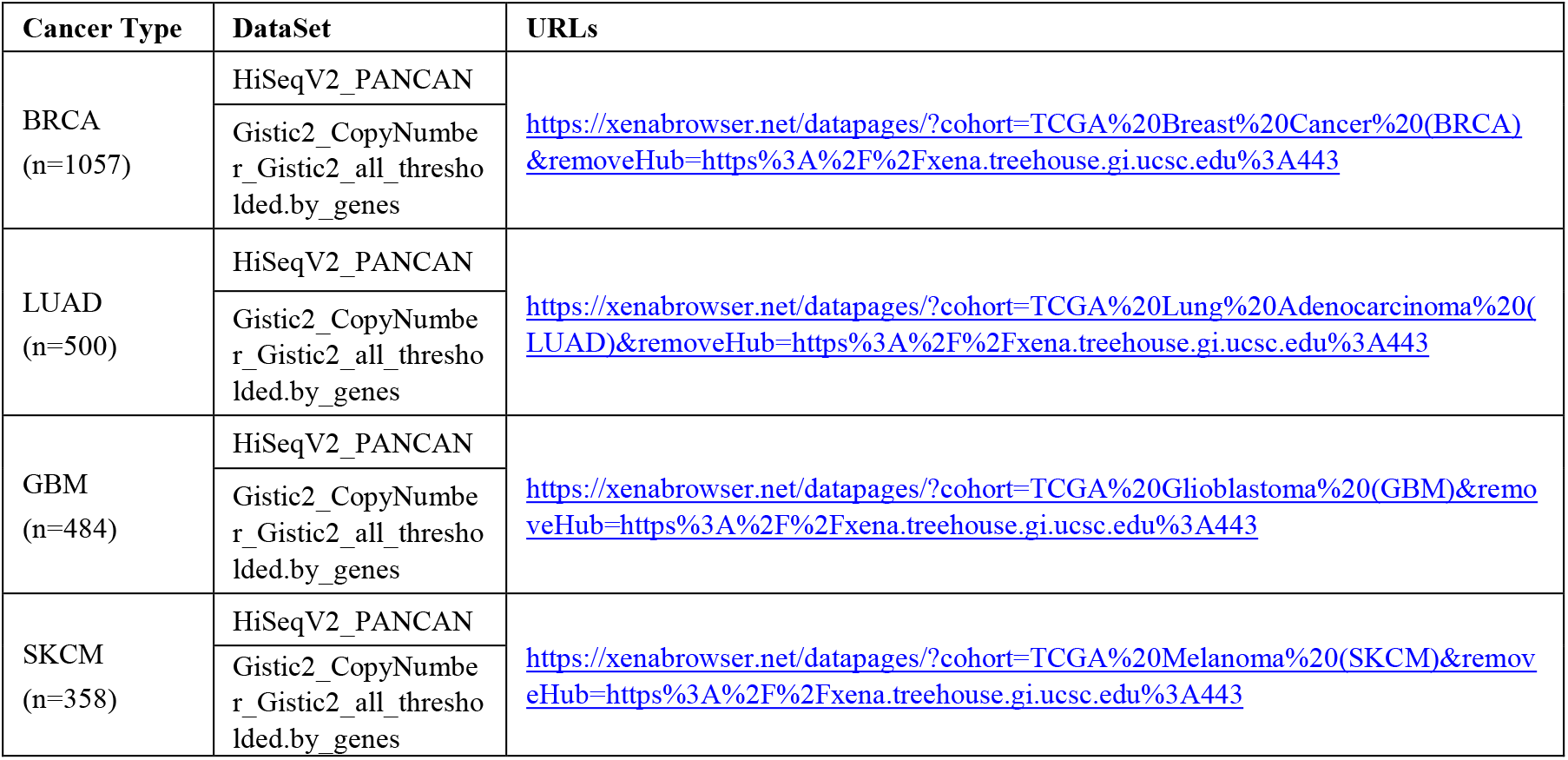
Number of samples, dataset_id and URLs to download the gene expression and copy number data from UCSC Xena data server.

### The 46 major signaling pathways

KEGG (Kyoto Encyclopedia of Genes and Genomes)^10^ is a database for the systematic understanding of gene functions. The KEGG signaling pathways provide the knowledge of signaling transduction and cellular processes. There are 303 pathways in KEGG database, and 45 of them are annotated as “signaling pathways”. Many of the signaling pathways are important oncogenic signaling pathways^9^, e.g., EGFR, WNT, Hippo, Notch, PI3K-Akt, RAS, TGFβ, p53. The ‘cell cycle’ cellular process is also included. For simplification, the ‘cell cycle’ is also viewed as one ‘signaling’ pathway. In total, 46 signaling pathways (45 signaling pathways + cell cycle) are selected (see **Table II**). Among these 46 signaling pathways, there are 1648 genes with both gene expression and copy number variation data. In summary, there are gene expression (TPM) and copy number variation data of 1648 genes in 46 signaling pathways of 45 cancer cell lines, which was used as the input of the deep learning model.

**Table II:** The 46 signaling pathways used for the analysis.

|  |  |  |  |  |
| --- | --- | --- | --- | --- |
| MAPK | FoxO | TGF-beta | T cell receptor | Adipocytokine |
| ErbB | Sphingolipid | VEGF | B cell receptor | Oxytocin |
| Ras | Phospholipase D | Apelin | Fc epsilon RI | Glucagon |
| Rap1 | p53 | Hippo | TNF | Relaxin |
| Calcium | mTOR | Toll-like receptor | Neurotrophin | AGE-RAGE |
| cGMP-PKG | PI3K-Akt | NOD-like receptor | Insulin | Cell cycle |
| cAMP | AMPK | RIG-I-like receptor | GnRH |  |
| Chemokine | Wnt | C-type lectin receptor | Estrogen |  |
| NF-kappa B | Notch | JAK-STAT | Prolactin |  |
| HIF-1 | Hedgehog | IL-17 | Thyroid hormone |  |

### Model Architecture of the DeepSigSurvNet

**Fig. 1** shows the schematic architecture of the proposed *DeepSigSurvNet* model. In the ‘input layer’, there are 2 input features, i.e., normalized gene expression across TCGA samples and integer copy number, for each of 1648 genes. Then the genes are connected to the 46 signaling pathways, only if a gene is included in a signaling pathway (not fully connection). The gene connection matrix and pathway connection matrix are used for specific the connections. The output of the 46 signaling pathways will be used as the input the convolution and inception layers (see **Fig. 1**). The activation functions for the Dense and convolution layers are the ReLu activation function. The last dense layer uses a linear activation function. To better model and predict the survival time of cancer patients, 3 clinical factors: age, gender and stage and the vital status are concatenated with the genomics data. For the training parameters, the batch size is 32, optimizer is “adam”. We divided the cancer samples in each type of cancer into training (80%) and test data (20%). For four cancer type, we use the same model architecture with different dropout rate, regularization value, and epoch. To investigate the relevance of individual signaling pathways for the survival time prediction, we employed the Layer-Wise Relevance Propagation (LRP) approach, which is available in the “iNNvestigate” package^13^. The distributions of the relevance scores, estimated by using the kernel density estimation based on the relevance scores of all samples, of all 46 signaling pathways for each type of cancer are obtained to investigate and understand the relevance of individual signaling pathways to the patients’ survival.

**Figure 1:**
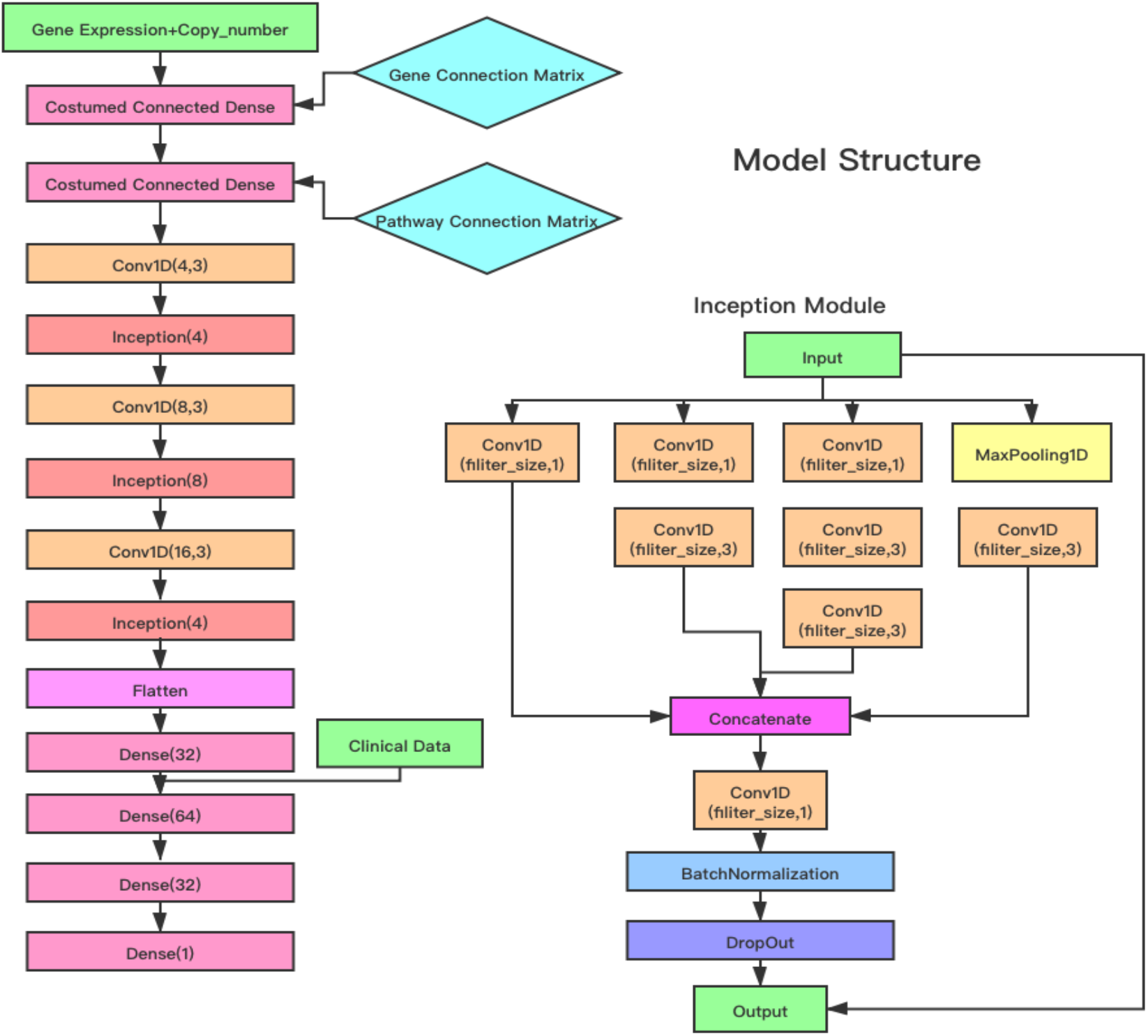
Schematic architecture of the DeepSigSurvNet model.

## Results

### Model performance evaluation

To evaluate the performance of the proposed model, the concordance index (c-index)metric was used. The c-index is defined as follows. Let *y*_*i*_ and *ŷ*_*l*_ be the true and predicted survival time. The concordance is defined as *P*(*ŷ*_*l*_ > *ŷ*_*j*_|*y*_*i*_ > *y*_*j*_), where *i* and *j* are two randomly selected samples. The c-index indicates the probability that prediction and the real survival time are relatively consistent or concordant, i.e., *ŷ*_*l*_ > *ŷ*_*j*_, *and y*_*i*_ > *y*_*j*_, or *ŷ*_*l*_ < *ŷ*_*j*_, *and y*_*i*_ < *y*_*j*_. Let C, D, T represent the number of concordant, discordant, and equal survival time, then the c-index is defined as:

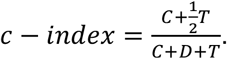

We compared the proposed model with random forest model, which is available as in the RandomForestRegression from the scikit-learn package. We train the random forest model using the same training and test dataset setting for the four types of cancer. The “n_estimator” and “max_depth” parameters are used to find the best performance of the random forest models. For the DeepSigSurvNet model, we use same architecture for all four types of cancer, with different dropout rate, regularization value and epoch number for each cancer type. **Table III** and **Table IV** show the comparison results. As can be seen, the random forest model has the similar c-index values in the training datasets. However, it has much lower c-index values on the test datasets, compared with the proposed DeepSignSurvNet model, which indicates that the proposed deep learning model is robust.

**Table III:** C-index values of random forest model in four types of cancer.

| Data set | n_estimator | Max_depth | c-index |
| --- | --- | --- | --- |
| Training-GBM | 30 | 5 | 0.6550 |
| Test-GBM | 30 | 5 | 0.5598 |
| Training-BRCA | 40 | 7 | 0.7849 |
| Test-BRCA | 40 | 7 | 0.5946 |
| Training-LUAD | 30 | 6 | 0.7433 |
| Test-LUAD | 30 | 6 | 0.5593 |
| Training-SKCM | 60 | 9 | 0.9419 |
| Test-SKCM | 60 | 9 | 0.5112 |

**Table IV:**
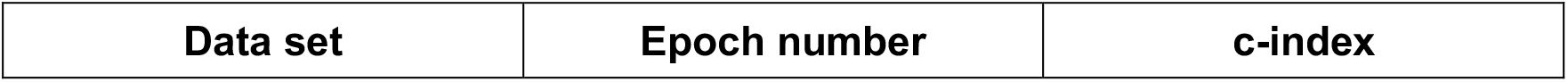
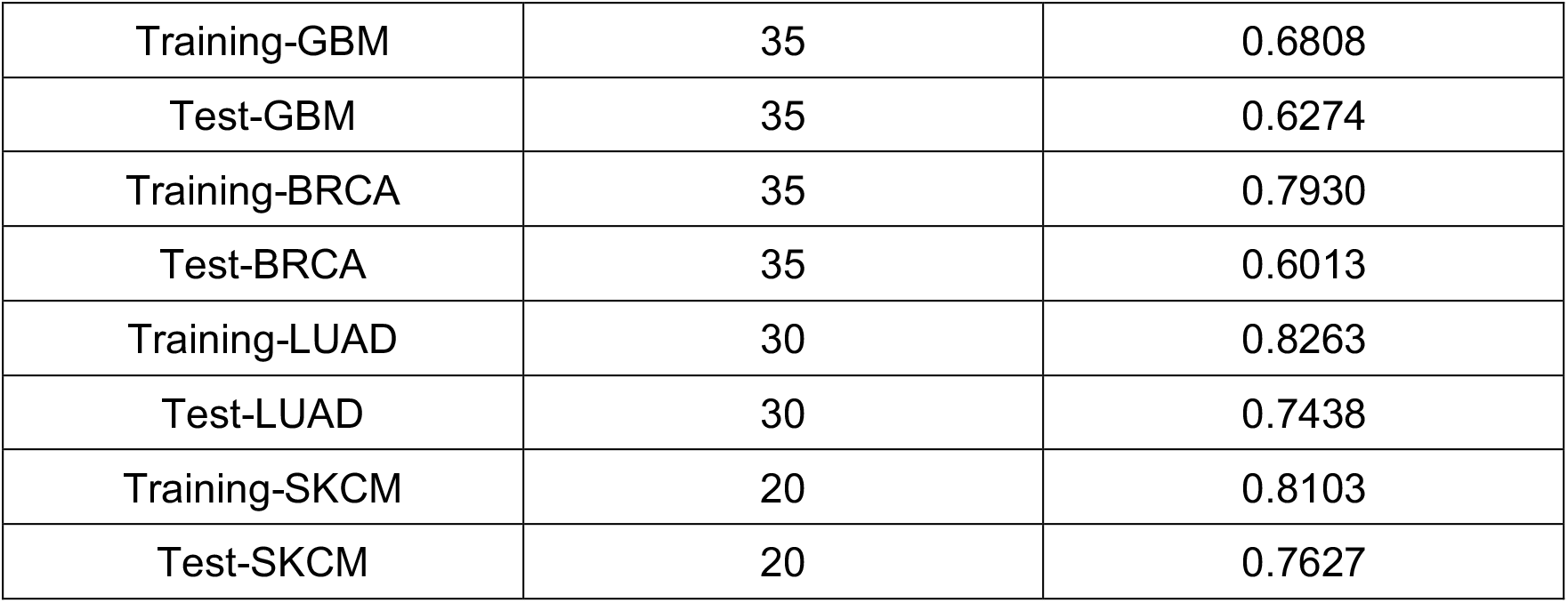
C-index values of DeepSigSurvNet in four types of cancer.

### Relevance of individual signaling pathways in 4 types of cancer

As discussed, it is interested to investigate and understand how the individual signaling pathways contribute to the cancer patients’ survival prediction. After training the deep learning models, we employed the ‘iNNvestigate’ package to calculate the relevance scores of the individual signaling pathways on individual cancer patients in each of 4 types of cancer. **Fig. 2** and **Fig. 3** show the probability density distributions of 46 signaling pathways in the 4 types of cancer.

**Figure 2:**
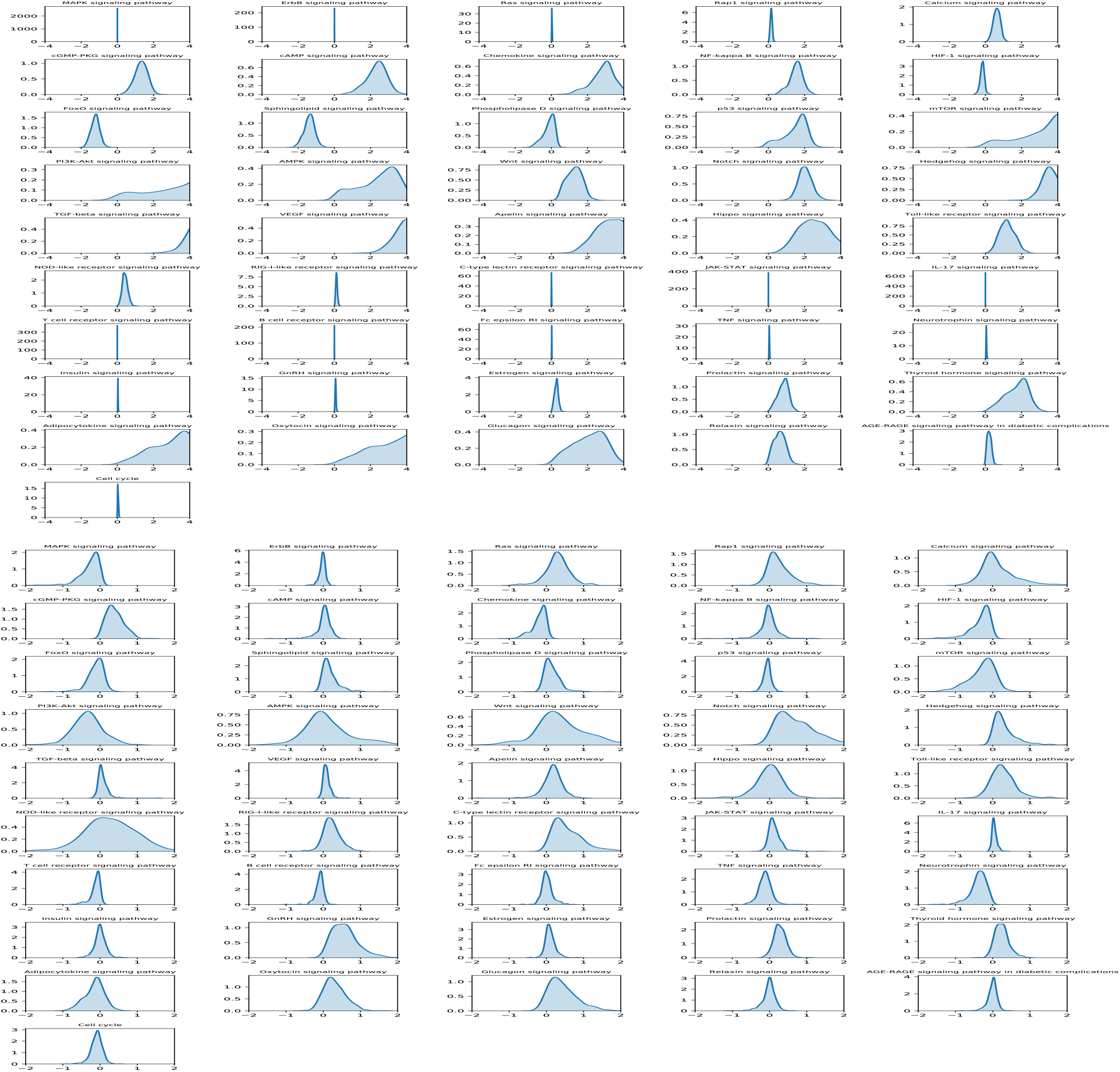
Density distribution of relevance scores of 46 signaling pathways on BRCA (top) and LUAD (bottom) cancer.

**Figure 3:**
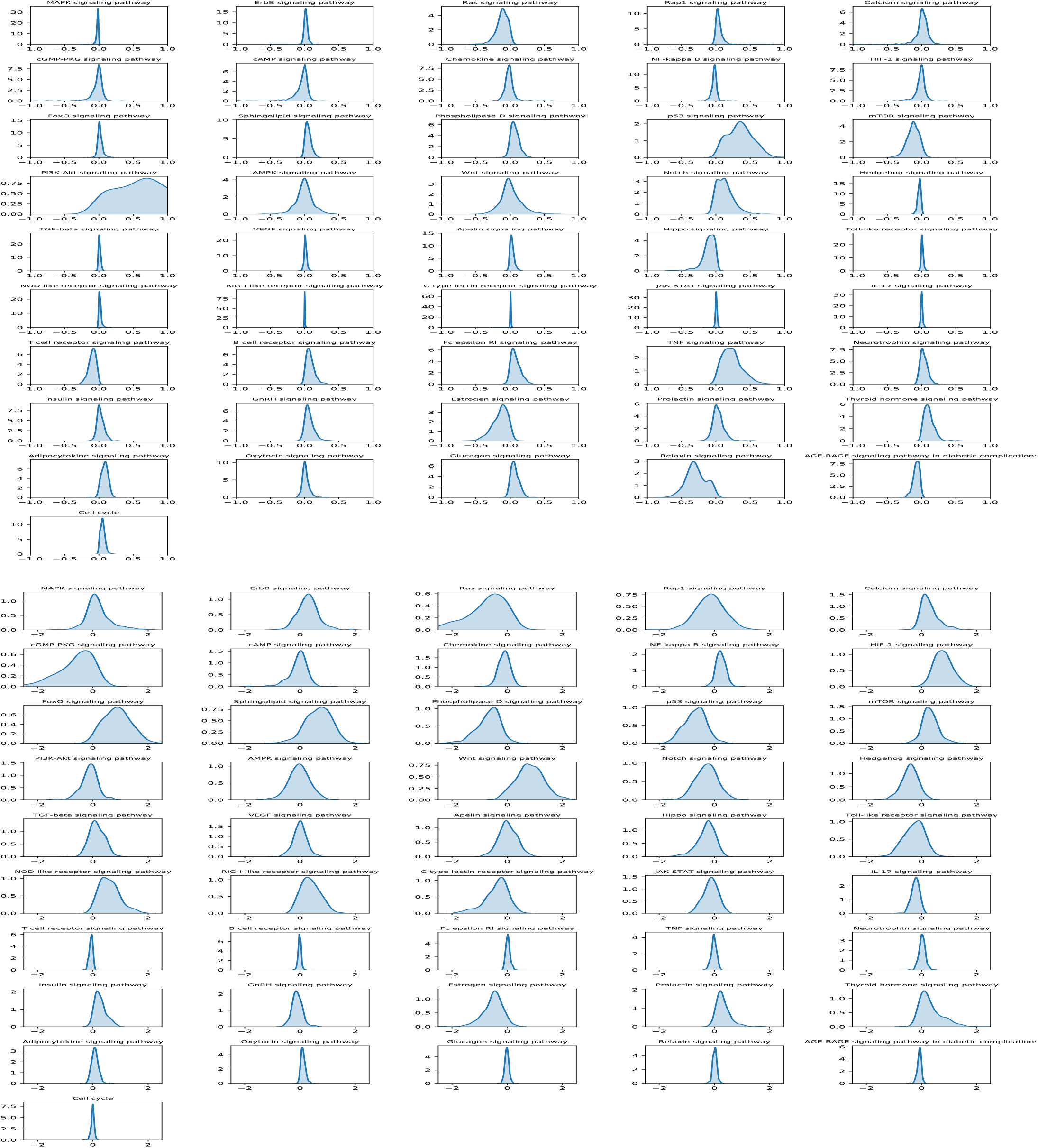
Density distribution of relevance scores of 46 signaling pathways on GBM (top) and SKCM (bottom) cancer.

Specifically, for BRCA, the mTOR, Hedgehog, PI3K-Akt, TGF-beta, AMPK, VEGF, Apelin, Adipocytokine and Oxytocin signaling pathways have the strongest relevance scores. P53, Wnt, Notch, NF-Kaapa B, FoxO, cGMP-PKG, cAMP, Chemokine, Sphingolipid, Relaxin, Thyroid hormone signaling pathways have relative high relevance scores. Surprisingly, the MAPK, ErbB, Ras, Rap1, JAK-STAT signaling pathways as well as cell cycle are not well associated with patients’ survival outcome. It is well known that these signaling pathways play important roles in cancer development. However, they can be separable in BRCA cancer samples to be the essential signaling pathways for patients’ survival outcome prediction. For LUDA, the patterns of density distributions are different from BRCA. More signaling pathways show high but not very strong relevance scores. For example, the MAPK, Ras, Rap1, cGMP-PKG, HIF-1, mTOR, PI3K-Akt, Wnt, Notch Hedgehog, C-type lectin receptor, GnRH, Neurotrophin, and Thyroid hormone signaling pathways have relatively high and consistent relevance scores. On the other hand, the AMPK, Hippo and NOD-like signaling pathways have the zero-mean value but with great variance. Then it is hard to evaluate their relevance important in cancer patients’ survival prediction analysis. For GBM, the Ras, p53, mTOR, PI3K-Akt, Notch, Hippo, TNF, Estrogen, Thyroid hormone and Relaxin signaling pathways have relatively high relevance scores; and the other signaling pathways are not correlated to the patients’ survival. For the SKCM, the patterns are kind of similar as the LUAD cancer samples. The Ras, Calcium, cGMP-PKG, NF-Kappa B, HIF-1, FoxO, Sphingolipid, Phospholipase D, p53, mTOR, Wnt, Hedgehog, NOD-like receptor, Estrogen, Prolactin, Thyroid hormone signaling pathways have relatively high and consistent relevance scores. Whereas, the MAPK, Rap1, PI3K-Akt, AMPK, VEGF signaling pathways have zero-mean values but with great variance.

In summary, the probability density distribution patterns of all the 46 signaling pathways vary significantly among the 4 types of cancer. For example, the p53 and mTOR signaling pathways are strongly relevant to patients’ survival outcomes in BRCA, GBM, SKCM, but not in the LUDA cancer patients. The MAPK, RAS, Rap1, ErBB signaling pathways are the known important signaling pathways in cancer. However, they are not strongly correlated with cancer patients’ survival outcome in the prediction models. It might be because all of these important signaling pathways are always activated in cancer patients. Thus, they are important targets for cancer therapy, but not informative in terms of the survival time prediction. Also, the cell cycle signaling does not play an important role in the survival time prediction. Moreover, a small set of signaling pathways, e.g., T cell receptor, B cell receptor, Fc epsilon RI, TNF signaling pathways do not show important contributions to the survival of cancer patients across all 4 types of cancer. Also, for each type of cancer, the less than half of the signaling pathways have strong effects to the survival prediction. Thus, drugs and drug combinations that can inhibit these essential signaling pathways, and that can inhibit the signaling pathways with strong relevance scores for each type of cancer might be effective to improve cancer patients’ survival time and outcome.

## Discussion and conclusion

Survival prediction is important in cancer studies. Deep learning models also have been proposed for the survival prediction, and outperformed the classic Cox PH model. However, it is challenging to understand the contributions of individual genes considering the non-linear combinations of a large number of genomics features, e.g., gene expression, copy number variation. Signaling pathways are important in cancer research to understand the signaling cascades regulating cancer development and drug response. Instead of using a large number of genomics features, in this study, we proposed a relatively biologically meaningful and simplified deep learning model, DeepSigSurvNet, for the survival prediction. In the model, the gene expression and copy number data of 1648 genes from 46 major signaling pathways are used. The analysis of deep learning model on 4 types of cancer can identify the distinct patterns of these signaling pathways, which are helpful to understand the relevance of the signaling pathways in the context of survival analysis, and can be novel targets for drug and drug combination prediction to improve cancer patients’ survival outcome. In conclusion, the proposed deep learning model can correlate the signaling pathways (not individual genes level) with cancer patients’ survival time and outcome.

There are some limitations of the proposed model that need to be further addressed. In addition to the 46 signaling pathways, other KEGG pathways, like metabolism pathways, will be further evaluated. Moreover, Gene oncology^17^ (GO) terms provide alternative biologically meaningful biologically processes (BP) (gene sets). Also, other omics data, like protein, methylation, genetic mutation can be integrated conveniently to the model in addition to the copy number, gene expression data. As aforementioned, the important genes within the important signaling pathways can be used potential gene signatures to discover drugs using the connectivity map (CMAP)^15,16^. We will investigate these possible directions in the future work.

## Reference

1. Cox DR. Regression Models and Life-Tables. J R Stat Soc Ser B. 1972;34(2):187–202. doi:10.1111/j.2517-6161.1972.tb00899.x

2. Krizhevsky A, Sutskever I, Hinton GE. ImageNet classification with deep convolutional neural networks. In: Advances in Neural Information Processing Systems.; 2012.

3. Goodfellow I, Pouget-Abadie J, Mirza M, et al. Generative Adversarial Nets. In: Ghahramani Z, Welling M, Cortes C, Lawrence ND, Weinberger KQ, eds. Advances in Neural Information Processing Systems 27. Curran Associates, Inc.; 2014:2672–2680. http://papers.nips.cc/paper/5423-generative-adversarial-nets.pdf.

4. Rajkomar A, Oren E, Chen K, et al. Scalable and accurate deep learning with electronic health records. npj Digit Med. 2018;1(1):18. doi:10.1038/s41746-018-0029-1

5. Devlin J, Chang M-W, Lee K, Toutanova K. BERT: Pre-training of Deep Bidirectional Transformers for Language Understanding. J ArXiv. 2018:abs/1810.04805.

6. Chaudhary K, Poirion OB, Lu L, Garmire LX. Deep Learning-Based Multi-Omics Integration Robustly Predicts Survival in Liver Cancer. Clin Cancer Res. 2018;24(6):1248–1259. doi:10.1158/1078-0432.CCR-17-0853

7. Subramanian A, Tamayo P, Mootha VK, et al. Gene set enrichment analysis: A knowledge-based approach for interpreting genome-wide expression profiles. Proc Natl Acad Sci. 2005;102(43):15545–15550. doi:10.1073/pnas.0506580102

8. Mobadersany P, Yousefi S, Amgad M, et al. Predicting cancer outcomes from histology and genomics using convolutional networks. Proc Natl Acad Sci. 2018;115(13):E2970 LP–E2979. doi:10.1073/pnas.1717139115

9. Sanchez-Vega F, Mina M, Armenia J, et al. Oncogenic Signaling Pathways in The Cancer Genome Atlas. Cell. 2018;173(2):321–337.e10. doi:https://doi.org/10.1016/j.cell.2018.03.035

10. Ogata H, Goto S, Sato K, Fujibuchi W, Bono H, Kanehisa M. KEGG: Kyoto encyclopedia of genes and genomes. Nucleic Acids Res. 1999:28. doi:10.1093/nar/27.1.29

11. Bach S, Binder A, Montavon G, Klauschen F, Müller KR, Samek W. On pixel-wise explanations for non-linear classifier decisions by layer-wise relevance propagation. PLoS One. 2015. doi:10.1371/journal.pone.0130140

12. Li B, Dewey CN. RSEM: accurate transcript quantification from RNA-Seq data with or without a reference genome. BMC Bioinformatics. 2011;12(1):323. doi:10.1186/1471-2105-12-323

13. Maximilian Alber, Sebastian Lapuschkin, Philipp Seegerer, Miriam Hägele, Kristof T. Schütt, Grégoire Montavon, Wojciech Samek, Klaus-Robert Müller, Sven Dähne P-JK. iNNvestigate neural networks! In: ArXiv.; 2018.

14. Subramanian A, Tamayo P, Mootha VK, et al. Gene set enrichment analysis: A knowledge-based approach for interpreting genome-wide expression profiles. Proc Natl Acad Sci U S A. 2005;102(43):15545–15550. doi:10.1073/pnas.0506580102

15. Lamb J, Crawford ED, Peck D, et al. The connectivity map: Using gene-expression signatures to connect small molecules, genes, and disease. Science (80-). 2006;313(5795):1929–1935. doi:10.1126/science.1132939

16. Subramanian A, Narayan R, Corsello SM, et al. A Next Generation Connectivity Map: L1000 Platform and the First 1,000,000 Profiles. Cell. 2017;171(6):1437–1452. doi:10.1016/j.cell.2017.10.049

17. Gene Ontology Consortium T, Ashburner M, Ball CA, et al. Gene Ontology: tool for the unification of biology NIH Public Access Author Manuscript. Nat Genet. 2000;25(1):25–29. doi:10.1038/75556

